# Global phylogenomic assessment of *Leptoseris* and *Agaricia* reveals substantial undescribed diversity at mesophotic depths

**DOI:** 10.1101/2022.09.12.504660

**Authors:** JC Gijsbers, N Englebert, KE Prata, M Pichon, Z Dinesen, R Brunner, G Eyal, FL González-Zapata, SE Kahng, KRW Latijnhouwers, P Muir, VZ Radice, JA Sánchez, MJA Vermeij, O Hoegh-Guldberg, SJ Jacobs, P Bongaerts

## Abstract

Mesophotic coral communities are increasingly gaining attention for the unique biological species they host, exemplified by the numerous mesophotic fish species that continue to be discovered. In contrast, many of the photosynthetic scleractinian corals observed at mesophotic depths are assumed to be depth-generalists, with very few species characterised as mesophotic-specialists. This presumed lack of a specialised community remains largely untested, as phylogenetic studies rarely include mesophotic samples and have long suffered from resolution issues associated with traditional sequence markers. Here, we used reduced-representation genome sequencing to provide a phylogenomic assessment of the two dominant mesophotic genera of plating corals in the Indo-Pacific and Western Atlantic, respectively, *Leptoseris* and *Agaricia*. While these genome-wide phylogenies broadly corroborated the morphological taxonomy, they also exposed substantial undescribed diversity (at least 24 molecular clades across the 8 species), including deep divergences within genera and current taxonomic species. Five of the eight focal species consisted of at least two sympatrically-occurring, genetically distinct clades, which were consistently detected across different methods. The repeated observation of genetically divergent clades associated with mesophotic depths highlights that there are many more mesophotic-specialist coral species than currently acknowledged, and that an urgent assessment of this largely unstudied biological diversity is warranted.

## INTRODUCTION

Because mesophotic coral ecosystems (MCEs) occur at depths beyond the limits of regular SCUBA diving (~30-150 m depth), they remain relatively understudied compared to shallow coral reefs, despite equalling or even exceeding the area occupied by the latter [1]. Over the past decades, interest in these deeper coral reef communities has grown due to their potential to act as a refuge against disturbances (for species with large depth distributions) [2–5] and as habitats hosting unique biological communities [7–9]. Their uniqueness is exemplified by the diversity and continuous discovery of depth-specialist fish species at mesophotic depths [6,9], as well as the vast differences observed between shallow and mesophotic fish species assemblages [7]. Although a similar differentiation over depth has been observed for reef-building coral assemblages [7,10], only a few scleractinian coral species dominate the assemblage at lower mesophotic depths [2,11–15], and a small proportion of them are considered deep-specialists [15,16].

Visual assessments (*in situ* or through imagery) of mesophotic coral diversity are challenging due to the intricate scale of and significant intraspecific variation in morphological traits used to identify species [16,17]. Because collection-based assessments of coral diversity are rare for mesophotic reef corals (particularly for depths >~60 m) [10] and because reference collections contain mostly shallow water coral specimens [15,18], morphological differences among shallow and mesophotic species can easily go unnoticed. Moreover, genetic identification has been hampered by the lack of species-level resolution when using traditional sequencing markers in scleractinian corals [19–22]. Consequently, these methodological challenges have greatly hindered the ability to differentiate and identify putative new scleractinian coral species associated with mesophotic depths.

Throughout the tropics, mesophotic coral ecosystems host reef-building scleractinian coral species with predominantly plating growth forms, which maximize light capture by their symbiotic dinoflagellates to sustain photosynthesis in low light conditions at greater depths [23–25]. Most of these plating corals belong to the family Agariciidae, with the genera *Leptoseris* and *Pavona* dominating mesophotic coral communities in the Indo-Pacific, and *Agaricia* in the Western Atlantic [11,14,17,26]. In these genera, species can generally occur over wide depth ranges, cover large areas of substrate, and provide important habitat structure to other reef-associated mesophotic organisms [13,27,28]. The genus *Leptoseris* is particularly dominant at lower mesophotic depths (>60 m [24]), and has been reported at depths of 172 m [29]. In Eastern Australia (Great Barrier Reef and Western Coral Sea) it has been observed down to 125 m depth [30], with four taxonomic species –*Leptoseris scabra* [31], *L. glabra* ([18], *L. explanata sensu* [32]), *L. mycetoseroides* [33], and *L. hawaiiensis* [31]– observed to dominate scleractinian coral communities at mesophotic depths [14,15]. These species also have a wide distribution ranging from the Red Sea [26,34,35] to the Hawaiian Archipelago (albeit with narrower depth distributions; [17]). The Western Atlantic genus *Agaricia* comprises seven species, of which four are dominant members of the mesophotic coral communities [27,36,37]: *Agaricia lamarcki* [38], *A. fragilis* [39], *A. grahamae* [40], and *A. undata* [41]. These four species occur throughout most of the Caribbean basin, and *A. fragilis* even extends north to Bermuda [3] and south to the Brazilian coast [42]. The deepest *Agaricia* colony (*A. grahamae*) was reported from a depth of 119 m [43]. Many of these widespread *Leptoseris* and *Agaricia* species occur across large depth ranges, making them important candidates to address the question to what extent mesophotic coral communities harbour unique depth-specialized species.

Molecular assessments of the genera *Leptoseris* and *Agaricia* using traditional sequence markers have exposed polyphyletic patterns, often in discordance with morphology-based taxonomy [11,13,17,27,34], and frequently were unable to discriminate between some of the well-established and morphologically distinct agariciid species [11,19,20,22,44–47]. Nonetheless, despite the pervasive and well-known issues with these markers [21,22], the few molecular studies that have been undertaken on this critically important family of scleractinian corals (Agariciidae) have highlighted the potential for undescribed diversity and depth-differentiation [11,13,17,34]. Reduced-representation genome sequencing methods (e.g., sequencing of restriction site-associated DNA (RAD-seq) or ultra-conserved elements (UCEs)) have demonstrated their potential to overcome these issues, resulting in phylogenies that have strong support (e.g., [48–52]), providing a promising way forward to study the evolutionary relationships within the Agariciidae. The increased resolution of such reduced representation methods was demonstrated through recent population genomics studies of *Agaricia* species, revealing significant genetic structuring within all four species dominating Atlantic mesophotic communities [3,37,53,54]. Here, we build on these initial findings and present a phylogenomic assessment of the two dominant mesophotic genera found within the Agariciidae family (figure 1b-c) in the Indo-Pacific and Western Atlantic (figure 1a). Focusing on eight species from the genera *Leptoseris* and *Agaricia* (figure 1a-b), we evaluate whether current systematics give an accurate reflection of the species diversity and extent of specialisation (i.e., depth specificity) in mesophotic coral communities.

**Figure 1.**
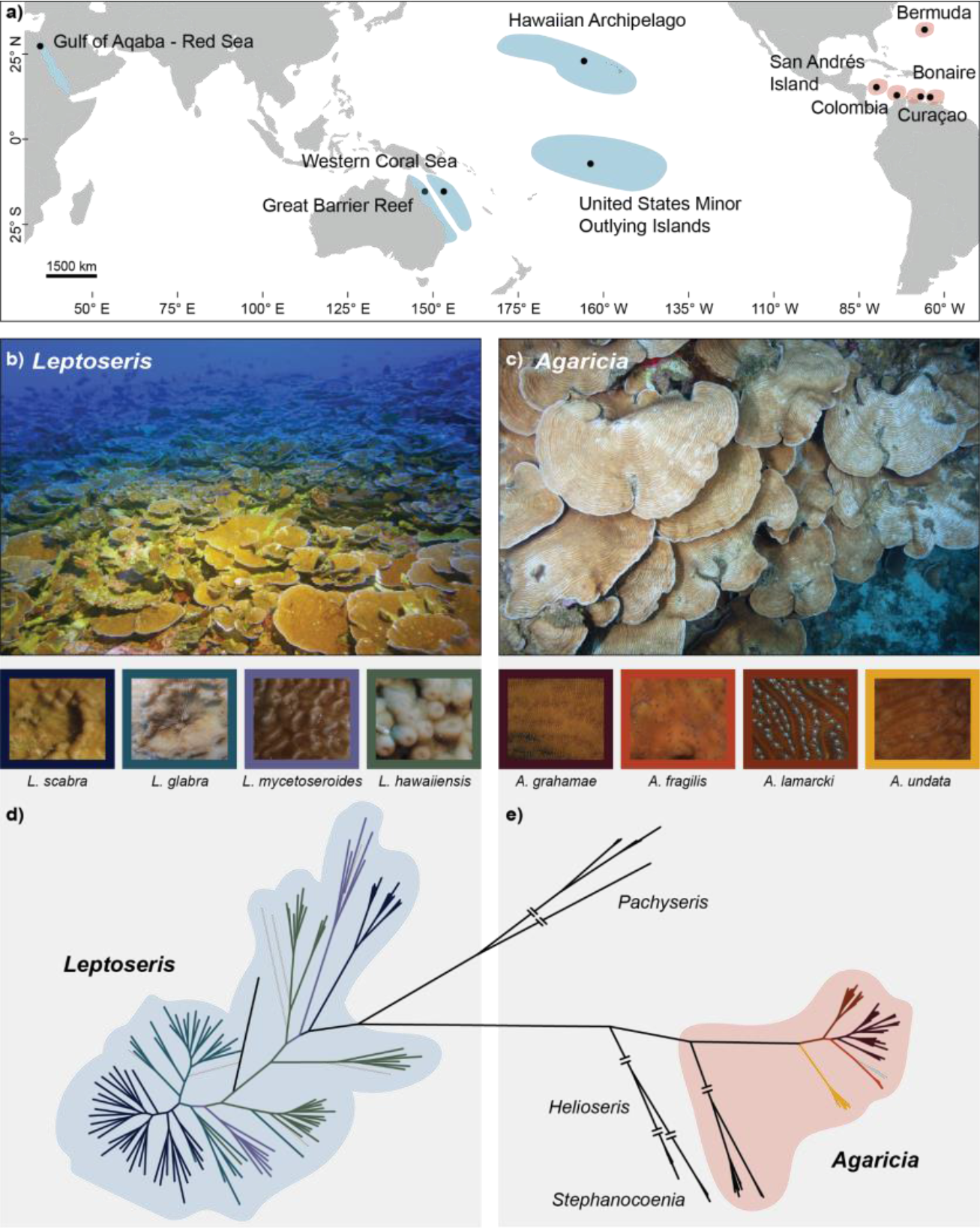
Overview of the *Leptoseris* and *Agaricia* study species. (a) Map of the sampling locations for *Leptoseris* in the Indo-Pacific (blue) and *Agaricia* in the Western Atlantic (orange). (b) *Leptoseris*-dominated coral community at 87 m depth in the Hawaiian Archipelago (Photo: Hawaiian Undersea Research Laboratory). (c) *Agaricia*-dominated coral community at 40 m in Curaçao, Southern Caribbean. (d) Phylogenetic tree (RAxML-ng) based on 37,528 concatenated nextRAD loci (3,361,114 sites) of the overall dataset, highlighting the position of *Leptoseris* (blue) and *Agaricia* (red) relative to included outgroups. Specimens from the focal species of this study are represented by coloured branches in the tree, with grey branches representing unidentified specimens.

## METHODS

### Sample collection and preparation

Coral specimens (n = 189) from the genera *Leptoseris* and *Agaricia* were collected as part of the “XL Catlin Seaview Survey” during visits to Eastern Australia (Western Coral Sea and Great Barrier Reef) and the Southern Caribbean (Curaçao and Bonaire; figure 1a, table S1). Samples were collected in Eastern Australia with a Seabotix vLBV300 Remotely Operated Vehicle [14,15] and in the Southern Caribbean with the “Curasub” submersible [11] and technical SCUBA. Small fragments (~1 cm^2^) were stored in 99% EtOH or NaCl 20% DMSO 0.5M EDTA solutions for genomic DNA extraction. Bleached skeletal specimens (all *Leptoseris*, and a subset of *Agaricia*) were deposited at the Queensland Museum Collection or the Invertebrate Zoology collection at the California Academy of Sciences (table S1). Additional tissue samples were acquired from collaborators in the Hawaiian Archipelago, US Minor Outlying Islands, Gulf of Aqaba (Red Sea), San Andrés Island, Cartagena, and Santa Marta (Colombia). The *Agaricia* dataset includes nextRAD sequence data (n = 52) from three published datasets [3,37,53], and we also included *Stephanocoenia intersepta* (n = 3 [3]), *Helioseris cucullata* (n = 5), and *Pachyseris speciosa* (n = 3 [55]) samples to serve as outgroups (table S1).

### Molecular dataset preparation

Genomic DNA was extracted as described in Bongaerts et al. [3,55], using the additional centrifugation steps to reduce endosymbiont contamination when sufficiently high gDNA yields were obtained. Extracted gDNA was used to create nextRAD DNA libraries (SNPsaurus, LLC). using selective PCR primers to genotype genomic loci consistently [56]. gDNA was fragmented and ligated with Nextera adapters (Illumina Inc). Once ligated, fragmented DNA was PCR-amplified (26 cycles, 73°C) with the matching primer for the adapter (“GTGTAGAGG”). Final libraries were sequenced (Illumina HiSeq 2500) to generate 100 bp single-end reads. Nextera adapters and low-quality ends (PHRED-quality score below 20) were trimmed using TrimGalore v0.6.4 (https://github.com/FelixKrueger/TrimGalore) to discard reads <30 bp and trim sequences up to 100 bp. IpyRAD v0.9.62 [57] was used for locus clustering and variant calling (85% clustering threshold, minimum coverage of six, minimum four samples per final locus, with all other settings run at default recommended values) (table S3).

Symbiont contamination was identified through a BLASTN comparison of each nextRAD locus against nextRAD sequence data from isolated Symbiodiniaceae, and four published Symbiodiniaceae genomes: *Symbiodinium microadriaticum* [58], *Breviolum minutum* [59], *Cladocopium goreaui* [60], and *Durusdinium trenchii* [61] removing positive matches (maximum E-value = 10^−15^) from the coral nextRAD loci. Potential microbial contamination was identified by an additional BLASTN comparison against the NCBI non-redundant database. The taxonomic IDs of positive matches (maximum E-value = 10^−4^) of non-cnidarian taxa were removed from the dataset. Three datasets were created from the filtered assembly and used for all downstream analyses: the *Leptoseris* dataset, composed of individuals from the genus *Leptoseris* with *Helioseris cucullata* and *A. fragilis* as outgroups (n = 127), the *Agaricia* dataset, composed of individuals from the genus *Agaricia* with *H. cucullata* and *L. glabra* as outgroups (n = 70), and the “*Agalepto*” dataset, composed of individuals from both genera with *H. cucullata*, *S. intersepta*, and *P. speciosa* as outgroups (n = 201) (figure 1d-e, table S1). NextRAD loci were then trimmed to 90 bp and filtered to retain loci genotyped for *ε*10 samples, with the resulting Variant Call Format (VCF) file filtered to retain only those SNPs genotyped for *ε*10% of samples.

### Phylogenetic and species tree inference

Maximum likelihood (ML) phylogenetic inference was performed using RAxML-ng v.1.0.1 [62] and a concatenated matrix with complete sequences of all loci. The best fit model of nucleotide substitution GTR+I+G4 was identified by ModelTest-ng x.y.z [63] and independent searches and bootstrap replicates were performed on each alignment until convergence was reached. To assess genealogical concordance, we used the concordance factor analysis as implemented in IQ-Tree 2.1.4-beta [64,65]. This analysis first infers single locus phylogenies (coupled with model selection) and then calculates the percentage of gene trees and alignment sites that are concordant with a reference ML topology (gCF and sCF, respectively). The reference ML topology was inferred in IQ-Tree using the concatenated matrix, applying an edge-linked proportional partition model [64] and employing standard model selection [66]. Species tree inference was performed using Tetrad, a coalescent species tree approach based on SVDquartets [67] implemented in IpyRAD v 0.9.65 [57]. This analysis constructs quartets using SNPs sampled from each locus and joins these to create a ‘supertree’ statistically consistent under the multispecies coalescent model. A single SNP was randomly sampled from each locus, all possible quartets were sampled, and 100 bootstrap replicates performed generating a majority-rule consensus tree and individual trees used to generate a density tree.

### Genetic structure using SNP-based analyses

To assess the genetic structure across our samples while avoiding the bias of *a priori* species assignment, we used *de novo* Discriminant Analysis of Principal Components (DAPC, [68]), an unsupervised dimensionality reduction method where novel genetic clusters are defined using a K-means clustering method and subsequently visualised using Principal Components Analysis (PCA). The DAPC analysis was conducted in R with the Adegenet package [69], choosing PCs based upon the optimal a-score and assessing the optimal numbers of clusters running K-means sequentially with increasing values of K, starting with K equal to the lowest AIC and BIC and then sequentially increasing K up to the maximum number of clades found in the ML tree.

### Comparison with traditional sequence markers

To compare the resolution of nextRAD with traditional mitochondrial markers, *cox*1-1-rRNA intron sequence data of 101 *Leptoseris*, 9 *Agaricia*, and 1 *Pavona* individuals were amplified using AGAH/AGAL primer pairs [70]. The PCR amplifications were performed following the approach of Bongaerts *et al*. [27]. Agarose gels were used to assess the quality of the PCR products, being then cleaned (ExoSAP-IT) and sequenced in forward and reverse directions (ABI BigDye Terminator chemistry, Australian Genome Research Facility). Additionally, previously published *cox*1-1-rRNA intron sequence data of 46 *Leptoseris* individuals [17], 61 *Agaricia* [11,71], 1 *Pavona clavus* [11], and 1 *L. hawaiiensis* [11] individuals were retrieved from GenBank. Codoncode Aligner was used to analyse the resulting sequences. ML phylogenies were inferred using RAxML-ng v.0.9.0 on the concatenated alignment, under the K80+G4 model, with a calculation of 20 trees and bootstrap support values based on 50,000 and 3,300 replicates for species from the genera *Leptoseris* and *Agaricia*, respectively.

## RESULTS

### Phylogenomic patterns across the genera *Leptoseris* and *Agaricia*

Using a reduced-representation sequencing approach (nextRAD), we recovered an average of 2.6 million reads (range: 580K-12.7M) from 201 scleractinian coral specimens; *Leptoseris* samples averaged 1.8 million reads (figure S1b), and *Agaricia* samples averaged 3.7 million reads (figure S2b). A maximum likelihood phylogenetic tree of the overall dataset, including outgroup genera (based on 37,528 nextRAD loci), confirmed that all of our ingroup specimens belonged to either the *Leptoseris* or *Agaricia* clade (figure 1d-e). One exception was a group of five presumed “*Leptoseris*” specimens from the Red Sea that grouped with the *Pachyseris* outgroup, and were later identified as *Pachyseris inatessa* (figure 1e). After separating the datasets and filtering, the *Leptoseris* dataset consisted of 15,250 nextRAD loci and 100,270 SNPs, and the *Agaricia* dataset of 19,902 nextRAD loci and 221,031 SNPs. For the genealogical concordance analysis, we inferred phylogenies for each single loci containing data for *ε*10% of the samples, resulting in 10,317 and 30,650 nextRAD loci for *Leptoseris* and *Agaricia*, respectively (figure S3c-d, figure S4c-d). The phylogenetic analyses supported the current taxonomic species within both genera but exposed substantial genetic substructure with at least 13 and 12 molecular clades observed within our focal *Leptoseris* and *Agaricia* species, respectively.

For the genus *Leptoseris*, maximum likelihood phylogenetic inference using both RAxML-ng and IQTree recovered extremely similar topologies consisting of many well-supported clades, particularly at deeper nodes (RAxML-ng: 6.92-100%, bootstrap range; IQTree: 35-100%; figure S3a-b). Gene (gCF) and site concordance factors (sCF) across the IQTree topology were lower and variable, particularly near the shallow nodes of the tree (gCF: 7.0 ± 12.2%, mean ± SD; sCF: 42.1 ± 16.8%; figure S3c-d), indicating the presence of discordant signal across loci and sites (most likely attributed to the short length of nextRAD loci). The species tree analysis using Tetrad recovered similar clades as those identified with ML, although the relationship among clades differs and their support varies (1-100%; 42.3 ± 30.4%; figure S5a). Clustering analysis (*de novo* DAPC) recovered clusters that corresponded with nodes at varying evolutionary depths in the ML and species tree topologies, with clusters identified at higher numbers of K corresponding with the same clades found in common among phylogenetic analyses (figure S6a). The exception was one group observed in a *L. glabra* clade (with mixed assignment to a spurious cluster that lacked individuals fully assigned to it).

While the clades recovered across methods for *Leptoseris* were largely composed of a single taxonomic species, all of the taxonomic focal species were represented by multiple clades. In several cases, these different clades were separated by relatively deep nodes (e.g. for *L. scabra* and *L. mycetoseroides*; figure 2a), whereas others represent substructuring within major clades (based on the tree topologies and the signatures of admixture at higher values of K from DAPC analysis; figure S6a). Samples belonging to the two divergent clades of *L. mycetoseroides* corresponded to respectively mesophotic samples from WCS (Western Coral Sea; 5 out of 6) and shallow water samples from Hawaii/USOMI (United States Minor Outlying Islands; 5 out of 6) (figure 2a). The former clade was identified as *L*. cf. *mycetoseroides* due to morphological variations from typical *L. mycetoseroides* (Pichon and Dinesen, personal observation). Similarly, while the two clades of *L. scabra* occurred sympatrically on reefs of the WCS and GBR (Great Barrier Reef), these clades were differentially characterised by depth, one clade was composed of a mix of shallow (10 – 20 m; n = 13) and mesophotic (40 – 60 m; n = 19) specimens and the other exclusively composed of mesophotic specimens (40 – 80 m, n = 14). There were three clades composed of *L. glabra* specimens; two had a wide geographic distribution (including Australia and Red Sea) and exclusively contain specimens from mesophotic depths (n = 4 and 6), and the third clade (middle *L. glabra* clade in figure 2a; also observed in Australia and Red Sea) represented a mix of shallow (10 – 20 m; n = 8) and mesophotic specimens (40 – 60 m, n = 10). *L. hawaiiensis* specimens were observed across four clades (figure 2a). One clade comprised samples from primarily lower mesophotic depths on the GBR and WCS, but matched a specimen identified as *Leptoseris* sp. 1 from Hawaii [17]. A second clade had widespread representation containing specimens from Australia, USOMI, and the Red Sea, and originating from depths down to 60 m. In contrast, the third *L. hawaiiensis* clade contained almost exclusively specimens from below 120 m depth in Hawaii, with a closely related individual originating from the GBR collected at 124 m depth. The fourth clade was identified as *L*. cf. *hawaiiensis* due to morphological variations from typical *L. hawaiiensis* (Pichon and Dinesen, personal observation), with samples from 60 m in the WCS. There were also five *Leptoseris* sp. that could not be identified further: one of them (87 m depth; WCS) was related to the former *L*. (cf.) *hawaiiensis* clades. Two samples (40 – 60 m depth; WCS) grouped within two distinct *L. glabra* clades. The remaining two *Leptoseris* sp. originating from shallow depths (9 and 22 m, Hawaii) grouped with the shallow *L. hawaiiensis* and *L. mycetoseroides* clades from Hawaii. The analyses also included a single specimen of *L. amitoriensis* from the Red Sea, which grouped closest to the clade containing *Leptoseris* sp. 1 (figure 2a).

**Figure 2.**
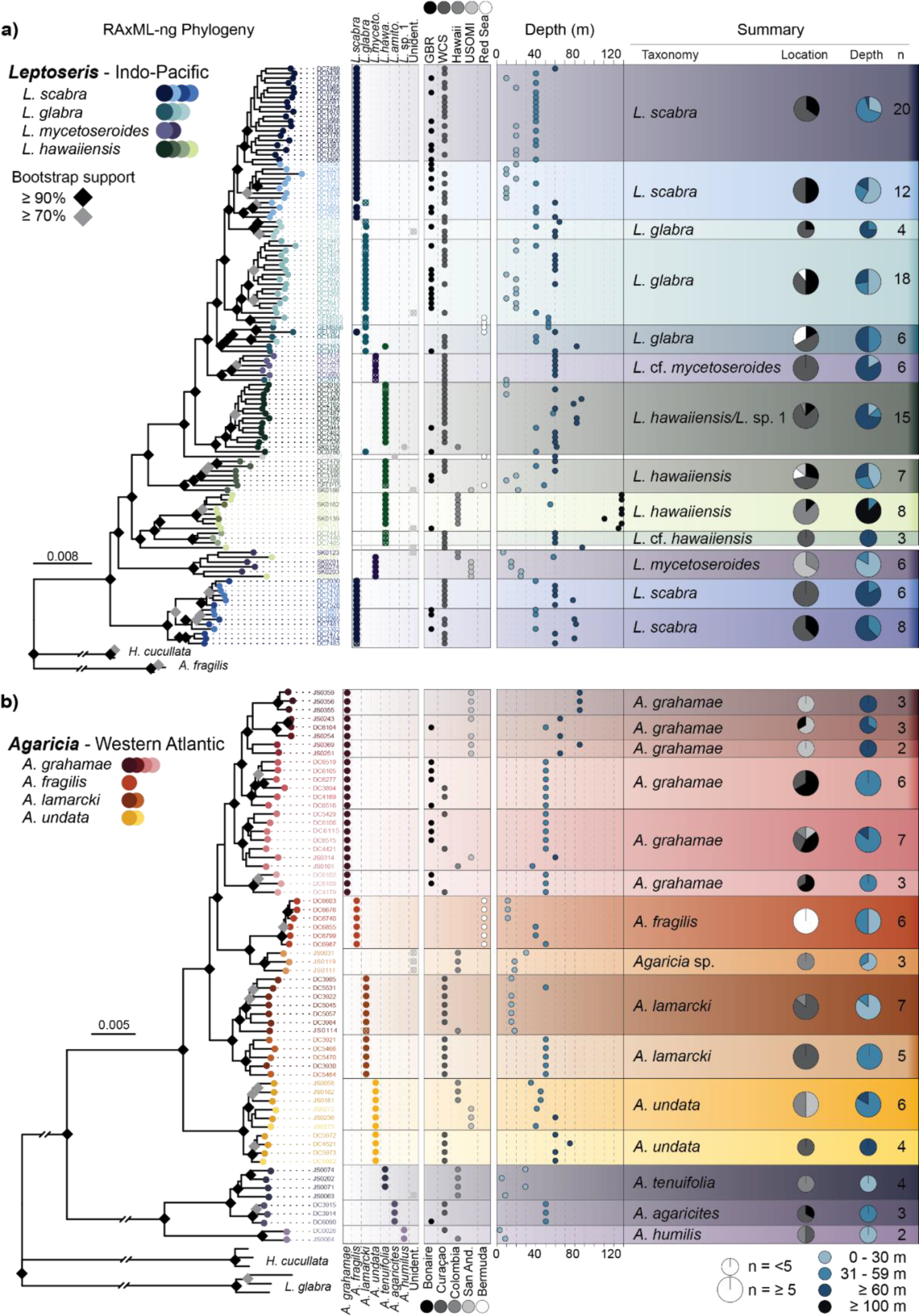
*Leptoseris* and *Agaricia*. (a) Phylogenetic tree (RAxML-ng) of the genus *Leptoseris* based on 15,250 concatenated nextRAD loci. (b) Phylogenetic tree (RAxML-ng) of the genus *Agaricia* based on 19,902 concatenated nextRAD loci. Colours across the trees represent the different identified clades (across phylogenetic and clustering methods) using blue, turquoise, purple, and green colours to represent the taxonomic species and shades of the different subclades. Columns next to the tree correspond to the taxonomic identification, sampling location, and sampling depth. The summary column on the right outlines the consensus taxonomic identification of each clade, and pie charts summarising the sampling depths and locations.

For the genus *Agaricia*, maximum likelihood phylogenetic inference using both RAxML-ng and IQTree also recovered very similar topologies consisting of several well-supported clades (RAxML-ng: 21.4 – 100%, bootstrap range; 80.3 ± 23.6%, mean ± SD; IQTree: 74 – 100%; 98.1 ± 5.3%; figure S4). Both phylogenies supported a major division of the currently acknowledged *Agaricia* genus into two major clades, representing the focal species, *A. grahamae*, *A. fragilis*, *A. lamarcki*, and *A. undata*, and the taxonomic species, *A. agaricites, A. humilis*, and *A. tenuifolia* [39] (figure 2b). Once again, the concordance factor values (gCF and sCF) across the IQTree topology were low and variable, particularly near the shallow nodes of the tree (gCF: 7.2 ± 12.4%, mean ± SD; sCF: 51.6 ± 22.2%; figure S4). The species tree analysis (Tetrad) recovered the same topology, consisting of the same genetic clades as those identified during ML analyses, with high statistical support, especially for the major clades (74 – 100%; 98.1 ± 5.3%; figure S5b). Clustering analysis (DAPC) recovered clusters that corresponded with nodes at varying evolutionary depths in the ML and species tree topologies, with the higher levels of K corresponding with the same clades found in the phylogenetic analyses (figure S6b).

Although each of the observed clades in the *Agaricia* phylogenies was composed of specimens of a single taxonomic species, all four taxonomic focal species were represented across two or more monophyletic subclades, with *A. grahamae* exhibiting further structuring into subclades that were supported by the DAPC clustering analyses (figure S6b). *A. grahamae* specimens were represented by four major sympatric clades (figure 2b). One clade (with substantial substructure) corresponded to the lower-mesophotic (>60 m) specimens from San Andrés (n = 7) and a single specimen from Bonaire upper-mesophotic reefs (figure 2b). The remaining *A. grahamae* clades consisted mostly of specimens from upper-mesophotic depths (<60 m) from Curaçao and Bonaire (n = 14), with one of those clades containing only two samples, one from Colombia and one from San Andrés. *A. lamarcki* consisted of two sympatrically occurring clades with one clade exclusively representing mesophotic specimens (n = 6) and the other mostly shallow water specimens (n = 6). The *A*. *undata* clade was further split into two clades corresponding to two sampling regions; the Southern Caribbean (Curaçao) containing lower-mesophotic specimens (n = 4) and South-western Caribbean (Colombia and San Andrés) consisting of upper-mesophotic specimens (n = 6), although with DAPC clustering showing admixture between clusters (figure S6b). Three specimens identified as *Agaricia* sp. from Colombia formed a separate clade grouping with *A. fragilis* from Bermuda (n = 6), with no further differentiation observed between shallow and upper mesophotic specimens of the latter. Specimens from *A. agaricites*, *A. tenuifolia*, and *A. humilis* formed a separate, highly divergent clade from the other *Agaricia* spp. Within this clade, *A. humilis* is the most divergent, with the *A. tenuifolia* from Colombia (n = 4) and *A. agaricites* from Curaçao and Bonaire (n = 3) forming closely related groups and exhibiting high levels of admixture in the DAPC clustering analyses (figure S6b).

### Comparison with traditional sequence markers

Phylogenetic analysis of the mitochondrial sequence marker *cox*1-1-rRNA using a ML approach (RAxML-ng) for a subset of the Australian *Leptoseris* specimens (n = 101, figure S7) including available sequences from Luck *et al*. [17], identified several major molecular clades with high bootstrap support that exceeded the number of focal taxonomic species (figure S7). Even though many of these clades consisted of specimens from a single taxonomic species, substantial polyphyly was observed with clades consisting of representatives from multiple taxonomic taxa. These clades do not appear to be described by geography nor bathymetric origins, with the exception of two divergent *L. scabra* clades consisting only of mesophotic representatives (1st: 40 – 101 m, n = 19, 2nd: 40 – 79 m, n = 7), as well as three *L. hawaiiensis* clades containing only *L. hawaiiensis* specimens mostly from mesophotic depths (1st: 60 m, n = 3; 2nd: 40 – 82 m, n = 8/9, 3rd: 40 – 60 m, n = 16, including an unidentified sample from 124 m, figure S7). For *Agaricia*, a comparison with available sequences (n = 61) from previous studies [11,71] and additional sequences (n = 9) using traditional sequence markers (*cox*1-1-rRNA) distinguished between three major clades, one consisting of *A. grahamae*, *A. fragilis*, *A. lamarcki*, one consisting of *A. undata*, and another one composed by *A. agaricites*, and *A. humilis*. However, this marker did not consistently discriminate between taxonomic species within these two groups (figure S8).

## DISCUSSION

Mesophotic coral ecosystems have gathered great scientific interest over the past decade, with numerous assessments evaluating the similarities and differences of these spatially extensive biological communities as compared to their shallow-water counterparts. From these, it has become clear that mesophotic communities become increasingly distinct with depth and can host unique and diverse species assemblages [7,15,72]. However, despite the growing number of accounts of depth-differentiated populations within scleractinian coral species (e.g., [3,73,74]), most scleractinian coral species at mesophotic depths are characterised as deep-generalists, with only a handful of reef-building scleractinian species recognised as mesophotic-specialists [16]. Through an extensive phylogenomic assessment of plating corals belonging to two dominant mesophotic genera in the Indo-Pacific and Caribbean (*Leptoseris* and *Agaricia*), we uncover substantial undescribed diversity with (1) assumed depth-generalists representing multiple depth-associated taxa, and (2) deep-specialist species consisting of multiple sympatric taxa. Overall, the results indicate that coral communities at mesophotic depths are more speciose and likely more specialised than currently acknowledged, urging for both systematic and ecological studies to capture and better understand this diversity.

### Phylogenomic insights into *Leptoseris* diversity

The observed phylogenomic patterns based on nextRAD data confirm that *Leptoseris* represent a taxonomically diverse group and comprise several highly divergent clades. When considering assignments based on the current *Leptoseris* taxonomy [18,32], species appear to be polyphyletic and often even separated by deep nodes (figure 2a), similar to previous studies using a mitochondrial intergenic spacer region [13,17,34]. Although the mitochondrial marker seemed initially promising for species delineation based on specimens solely from Hawaii [13,17], it did show pervasive polyphyly that extended across both *Leptoseris* and *Agaricia* genera [17], and a lack of genetic variation across morphologically divergent taxonomic species when applied in a different geographic region [34]. Using the same marker, we obtained similar results for specimens from Australia, where some of the taxonomic diversity is captured though not consistently, with taxonomic species spread widely across the tree (figure S7). In contrast, the reduced representation data shows much more consistent phylogenetic patterns, with all recovered clades consisting almost exclusively of a single taxonomic species (figure 2a). The patterns corroborate the current taxonomy based on morphological differences, but with the increased genomic resolution, sample sizes and geographic range, also unveiling additional diversity associated with specific geographic regions and bathymetric ranges. Given this congruence with taxonomic species and spatial distributions, the observed phylogenomic patterns are expected to more closely reflect the evolutionary relationships within this genus and demonstrate the resolving power of reduced representation methods.

Within the genus *Leptoseris*, the major split in *L. mycetoseroides* seems to correspond with both geography and depth (figure 2a). Given that the type specimen originates from the Marshall Islands (adjacent to USOMI) [75], there are no junior synonyms reported for this species [75], and several specimens from East Australia were noted as morphologically distinct, it is possible that the Hawaii/USOMI clade represents *L. mycetoseroides* with the mesophotic specimens in the East Australian clade representing an undescribed species. One of the clades identified for *L. hawaiiensis* consisted of predominantly lower mesophotic specimens from the WCS, and matched with a *Leptoseris* “sp. 1” specimen from Hawaii, where it was identified as a putative new species based on its distinct micromorphology [17]. In the *cox*1-1-rRNA phylogeny, one of our specimens from this clade also grouped with the *Leptoseris* “sp. 1” clade from Luck *et al*. [17], although the other specimens from the same nextRAD clade were widely spread across the tree (figure S7). This predominantly mesophotic clade is present in Australian waters and may be geographically widespread, although further investigations are warranted. The other three *L. hawaiiensis* clades each showed different depth distributions, with one from predominantly >100 m depth in Hawaii, and another representing morphologically atypical specimens from 60 m depth in Australia. Related to these clades but branching separately was a single specimen collected at the GBR at 124 m, representing the deepest published report of a zooxanthellate coral collected from the GBR [30], and indicating that *Leptoseris* clades occurring at the lower boundaries of mesophotic depths represent uniquely adapted species. We observed a similar partitioning into depth-associated clades for *L. glabra* and *L. scabra*, with a further subdivision into additional geographically sympatric clades for the latter. As reported by Luck *et al*. [17], the deeper clade of *L. scabra* is very divergent compared to the other *Leptoseris* taxa (although not resulting in generic polyphyly), and with *cox*1-1-rRNA sequences matching those from Hawaii. Detailed morphological characterization of the specimens (beyond identification according to currently acknowledged taxonomic species) was beyond the scope of the current study, but is the focus of ongoing work that aims to determine which of the exposed taxa are cryptic versus morphologically differentiated.

### Phylogenomic insights into *Agaricia* diversity

For the genus *Agaricia*, we observed a split into two major clades corresponding to species with deeper (*A. grahamae*, *A. fragilis*, *A. lamarcki*, *A. undata*) and shallower (*A. agaricites*, *A. humilis*, *A. tenuifolia*) distributions (figure 2b). This division was also observed using the *cox*1-1-rRNA region (figure S8) as well as other mitochondrial markers [11,27,71,76–78], although these markers have lacked the resolution to consistently discern the established taxonomic species within the clades. The division into two major clades is further corroborated by a split in gross morphology (unifacial versus bifacial colonies), microskeletal characteristics (including wall thickness and septocostae orientation; [79]), and genetic relatedness among symbiont associations within each major clade [27]. Based on a morphological analysis of modern and fossil representatives, Stemann [79] proposed these should represent two separate genera: *Agaricia* and *Undaria* [80]. Recent differences in spatial genetic structure observed across taxa further suggest major differences in reproductive strategies between these major clades [53]. Given the major phylogenetic divergence observed in this study (figure 2b), and further corroborated by the major locus dropout between clades (figure S2), we argue that the shallower and deeper *Agaricia* indeed represent two genetically, morphologically, and biologically divergent groups that might require generic reassignment.

The phylogenetic analyses supported the current taxonomic species within *Agaricia* but revealed substantial genetic substructure with at least 12 molecular clades observed within our four focal *Agaricia* species (figure 2b). This structure was consistently recovered across phylogenetic and clustering methods and often corresponded to different depths and/or geographic regions, indicating the role of both environmental and geographical (i.e., allopatric) contributors to the underlying diversification processes. Our analyses included four focal species previously assessed in population genomic studies with varying levels of intraspecific genetic structure [3,37,53]. Through a combined phylogenomic analysis, we were able to assess this genetic structuring in the context of interspecific variation. For example, despite the genome-wide differentiation observed for shallow and deep *A. fragilis* populations in Bermuda [3], representative specimens of those populations formed a single clade here (i.e., under a phylogenomic framework), indicating that these reflect the earlier stages of the divergence continuum. In contrast, the *A. lamarcki* samples clearly separated into two distinct clades associated with predominantly shallower (15 m) and upper mesophotic (50 m) depths (figure 2b) despite their geographically sympatric distribution and ongoing low levels of gene flow [53], and clearly represent distinct evolutionary units. Similarly, we observed four distinct clades of *A. grahamae* occurring sympatrically in Curaçao and Bonaire with varying levels of divergence (figure 2b), and while not partitioned by depth, it indicates undescribed mesophotic taxa that are currently not accounted for. The two clades observed for *Agaricia undata* corresponded with geography (Curaçao vs Colombia and San Andrés), whereas several Colombian specimens that could not be reliably identified down to species through morphological assessment (*Agaricia* sp. clade) grouped with *A. fragilis* from Bermuda, indicating the presence of this or an undescribed related species in the Southern Caribbean. In the aforementioned “*Undaria*” clade, *A. humilis* formed a separate clade from *A. agaricites* and *A. tenuifolia* specimens, corroborating their ecological and reproductive differences [36]. The separation between *A. agaricites* and *A. tenuifolia* was confounded by distinct geographic origins for the specimens; however, the lack of support in clustering analyses indicates that these morphologically similar species warrant further taxonomic investigation.

### Evolutionary patterns and remaining challenges

As a result of this study, we can now begin to directly compare and contrast the present diversity of these genera and to hypothesise the existence of associated generative processes. An evident pattern we observe across both genera is the depth-associated divergence within several focal species (e.g., *L. scabra*, *L. hawaiiensis*, or *A. lamarcki*). Depth is an important contributor to the divergence of marine species [73,81,82], with several well-studied examples in scleractinian corals (e.g., [83–87]). Our results corroborate the expectation that depth has led to divergent species associated with mesophotic depths [82], and that the underestimated diversity of mesophotic-specialist species is likely to be a consequence of logistical and taxonomic challenges [16]. However, depth does not immediately explain the sympatric taxa we observed within mesophotic-specialist species (e.g., *A. grahamae* and the “deep” *L. scabra* clade; lacking clear depth differences). Their ecological differentiation may be related to finer scale environmental conditions or, indeed, other factors and should be assessed within a spatially explicit framework targeted to microhabitat characterisation [88].

Another clear pattern we observe here is the higher levels of taxonomic/phylogenetic discordance in (often sympatrically-occurring) genetic groups in *Leptoseris*, relative to *Agaricia*. This difference might be the result of *Leptoseris* having a larger geographic distribution spanning a variety of ecoregions (Indo-Pacific and Caribbean; [89,90]) in contrast to the relatively restricted geographic distribution of *Agaricia* in the Western Atlantic and Caribbean [89]. In addition, the fossil record of *Leptoseris* dates back to the Oligocene (~23 Ma years, [91]), while *Agaricia* is a younger genus, dating from the Neogene ~12 Ma years [92], and likely diversified over the last ~3 Ma years following the closure of the Central American Seaway [89,92]. These differences suggest increased opportunities for ecological diversification in *Leptoseris* (compared to *Agaricia*), while signatures of admixture (figure S6a) suggest a possible history of hybridization and reticulation (i.e., the process of genetic lineages both merging and/or diverging through time; [93]) among its lineages. It is important to note, however, that the number of loci and sites supporting some of these polyphyletic patterns are low (figure S3c), thus additional sequence data (e.g., longer/more informative loci, whole-genome data) might reveal additional patterns within this genus (see below).

The reduced representation sequencing (nextRAD) data across multiple analytical approaches, consistently recovered groupings of individuals (i.e., clades and/or clusters) coherent with biological, ecological, or morphological evidence. Compared to traditional sequence markers, our genome-wide sequencing approach resulted in a higher resolving power, confirming the status of current morphologically and ecologically divergent taxonomic species, and allowing us to gain a better insight into the evolutionary history of members of both genera. Despite the consistent topology recovered across approaches, the support for these underlying phylogenies did vary across analytical methods. For example, despite the strong bootstrap support recovered across all ML phylogenies, the gene- and site-concordance analysis revealed variable values in both datasets, indicating, on average, a low fraction of gene trees and a moderate fraction of alignment sites supporting each topology (figure S3 and figure S4). This discrepancy between methods is often observed across different RAD-seq approaches, in most cases, due to the short length of each locus and its associated phylogenetic information [94–97]. Targeted sequence-capture approaches focused on ultra-conserved elements, or exons, can potentially represent an alternative to recover greater concordance across loci. Because conserved regions are less affected by demographic events, and next to these regions, more variable sites can be considered side by side, these genomic regions provide a more holistic view of individual evolutionary histories [98–99]. Compared to RAD-seq, however, the generally greater cost of such sequence-capture approaches often translates to smaller sample sizes and less replication. Ultimately, whole-genome resequencing of representatives across a wide range of depths and geographies from our study will enable us to identify the loci involved in the diversification of *Leptoseris* and *Agaricia* and help assess divergence histories through introgression events.

## Conclusions

Our results shed light on the diversity of two key genera of mesophotic ecosystems, *Leptoseris* and *Agaricia*. Using genome-wide sequencing data in a phylogenomic framework, we observed that genomic data corroborate current morpho-taxonomic criteria, but also exposed substantial undescribed diversity associated with mesophotic depths. Distinguishing where these different taxa sit along the speciation continuum remains challenging, particularly as traditional species delimitation methods are known to oversplit when using genomic data [100]. Nonetheless, current taxonomic species were observed to comprise highly divergent clades (e.g., *L. scabra*, *L. mycetoseroides*) or sympatrically occurring but geographically widespread subclades (e.g., *L. glabra*, *A. grahamae*) across phylogenetic and clustering methods, indicating that a reasonable extent of reproductive isolation has evolved and that most of these clades represent distinct species. Further integrative taxonomic studies are currently being developed to formally describe the uncovered species diversity, and verify whether these taxa are morphologically cryptic, differentiated and/or potentially align with junior synonyms. Overall, our study highlights how our perception of mesophotic coral ecosystems is affected by our shallow knowledge bias, and that studying the ecology and evolution of this newly exposed mesophotic biodiversity should be a priority in order to advance our understanding of these ecosystems.

## Supporting information

Supplementary figures Gijsbers et al 2022

## Acknowledgements

We thank for logistical support: CARMABI Research Station, Substation Curaçao, CIEE Bonaire Research Station, STINAPA Bonaire, CORALINA, PNN “Corales de Profundidad”, Hawaii Undersea Research Laboratory (HURL), Reef Connections, Mike Ball Dive Expeditions, SY Ethereal, and Waitt Foundation.

