## Supplementary figures Gijsbers et al 2022 for "Global phylogenomic assessment of *Leptoseris* and *Agaricia* reveals substantial undescribed diversity at mesophotic depths"

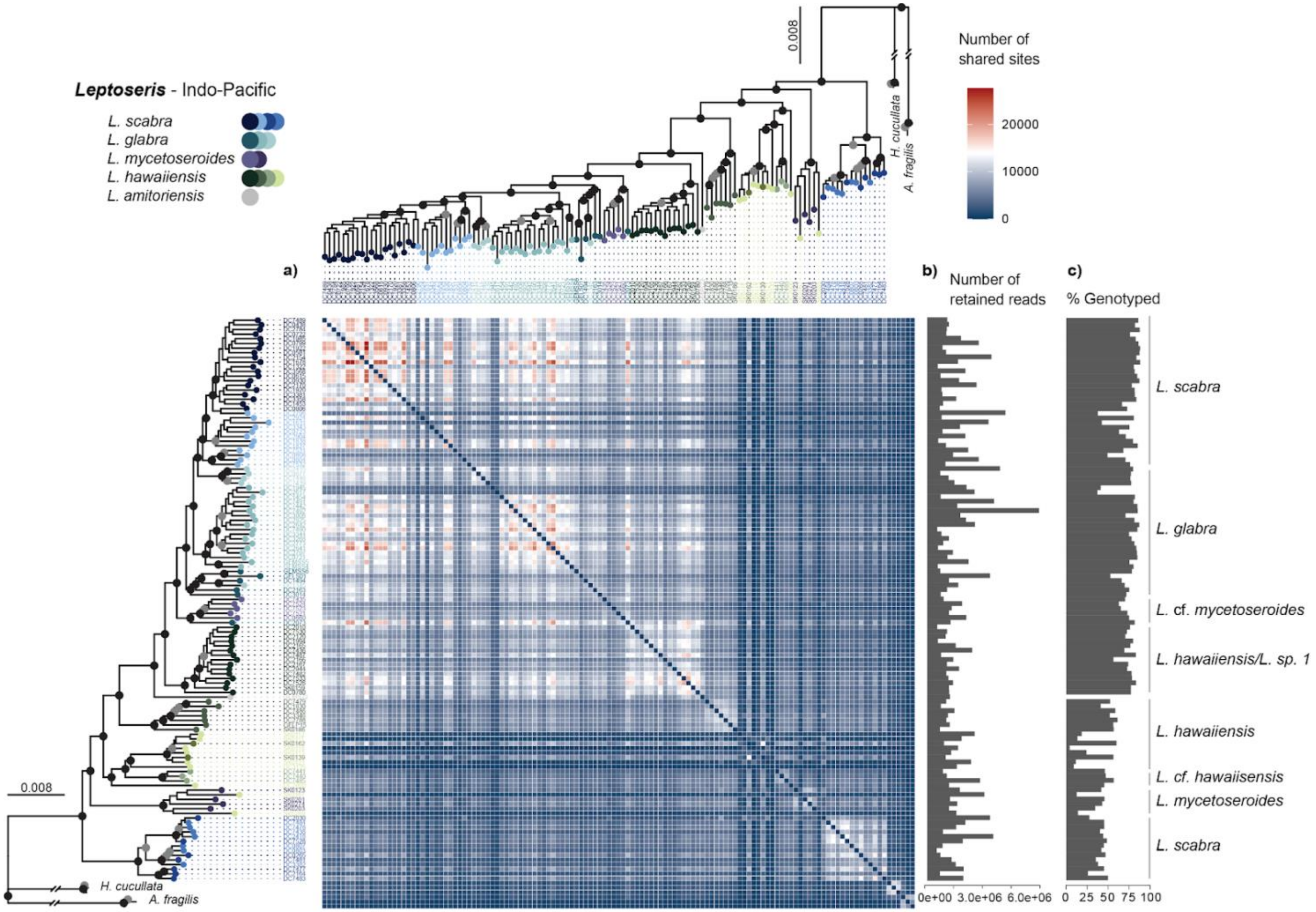


**Supplementary figure S1. Number of shared sites among *Leptoseris* specimens.** (a) Heat map representing the number of shared sites between individuals. Red and Blue correspond respectively, to the highest and lowest amount of shared sites between individuals. (b) Barplot representing the number of retained reads across individuals. (c) Barplot representing the percentage of genotyping across individuals.

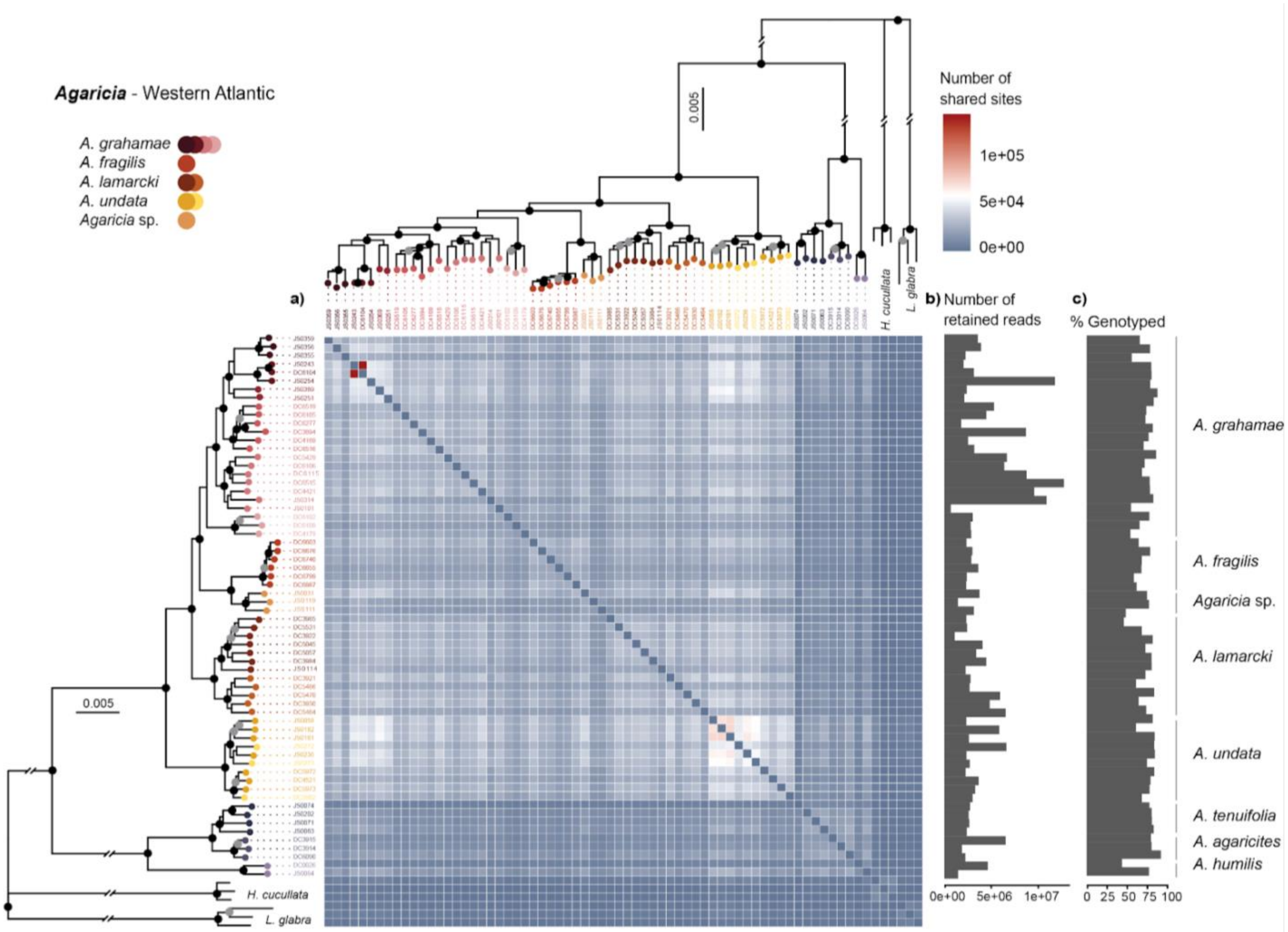


**Supplementary figure S2. Number of shared sites among *Agaricia* specimens.** (a) Heat map representing the number of shared sites between individuals. Red and Blue correspond respectively, to the highest and lowest amount of shared sites between individuals. (b) Barplot representing the number of retained reads across individuals. (c) Barplot representing the percentage of genotyping across individuals.

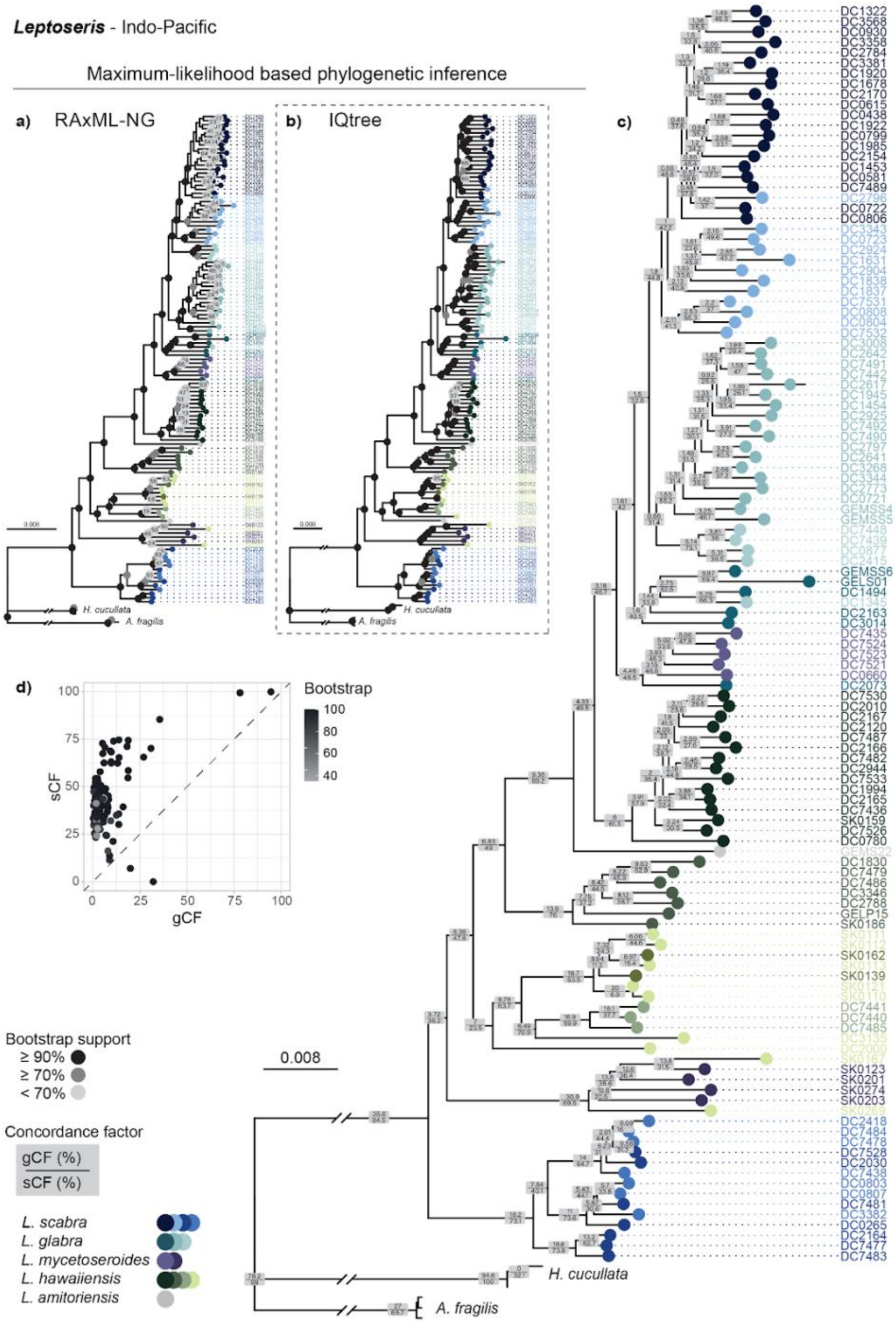


**Supplementary figure S3. Genealogical concordance for the *Leptoseris* dataset.** (a) RAxML-NG tree based on 15,250 concatenated nextRAD loci. (b) IQtree species tree based on 10,317 single full loci. (c) IQtree tree from inset with concordance factor values. Numbers in grey squares on each branch represent the gene concordance factor (gCF) above and the site concordance factor (sCF) on the bottom. (d) Scatter plot of sCF against gCF values in relation to the bootstrap values of 1000 ultrafast bootstrap replicates.

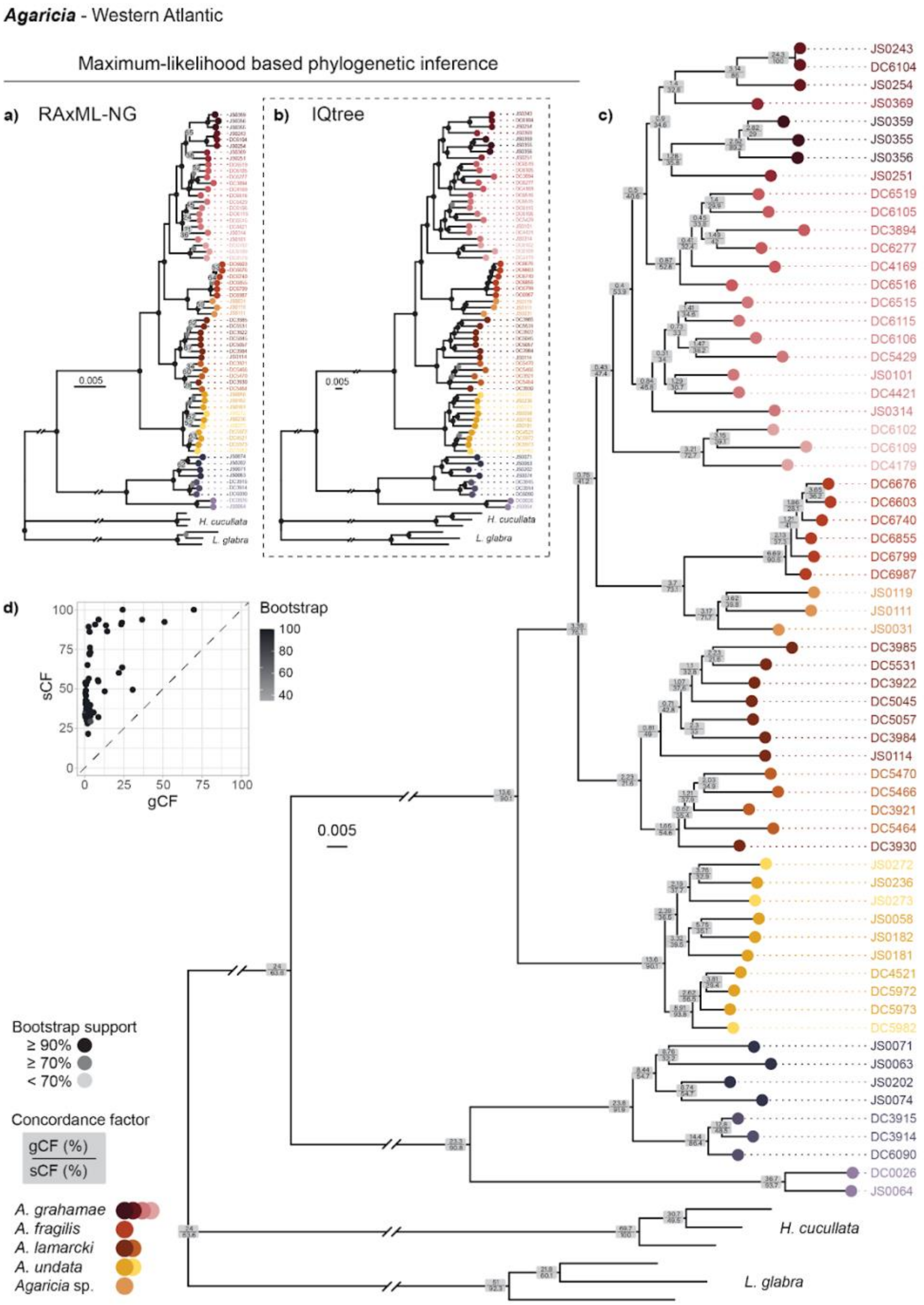


**Supplementary figure S4. Genealogical concordance for the *Agaricia* dataset.** (a) RAxML-NG tree based on 19,902 concatenated nextRAD loci. (b) IQtree species tree based on 30,650 single full loci. (c) IQtree tree from inset with concordance factor values. Numbers in grey squares on each branch represent the gene concordance factor (gCF) above, and the site concordance factor (sCF) on the bottom. (d) Scatter plot of sCF against gCF values in relation to the bootstrap values of 1000 ultrafast bootstrap replicates.

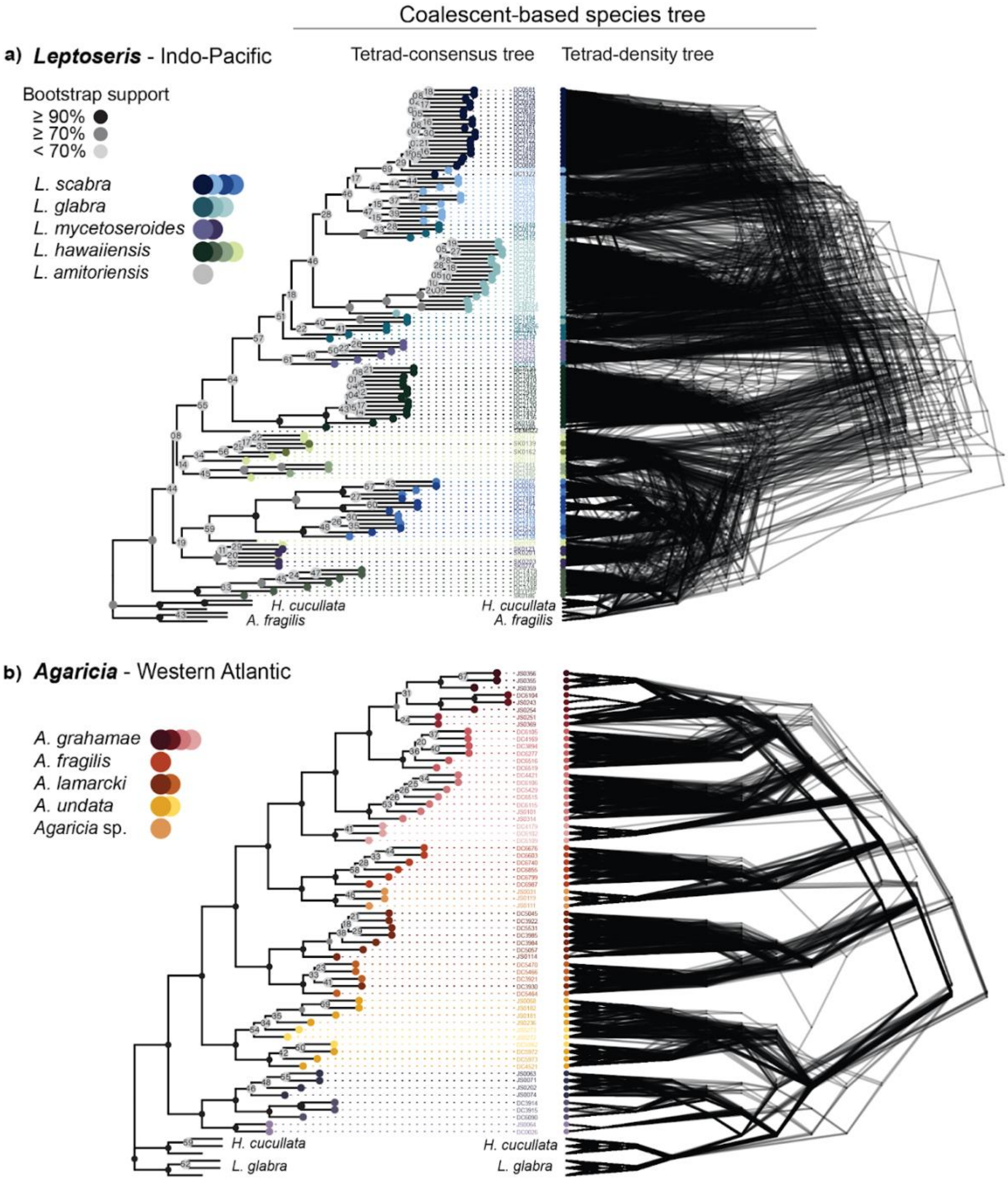


**Supplementary figure S5. Evolutionary relationships of mesophotic *Leptoseris* and *Agaricia* species.** (a) *Leptoseris*: Coalescent-based species tree: Tetrad; consensus tree of 100 bootstrap replicates (left), density tree visualizing 100 bootstrap trees (right). (b) *Agaricia*: Coalescent-based species tree: Tetrad; consensus tree of 100 bootstrap replicates (left), density tree visualizing 100 bootstrap trees (right).

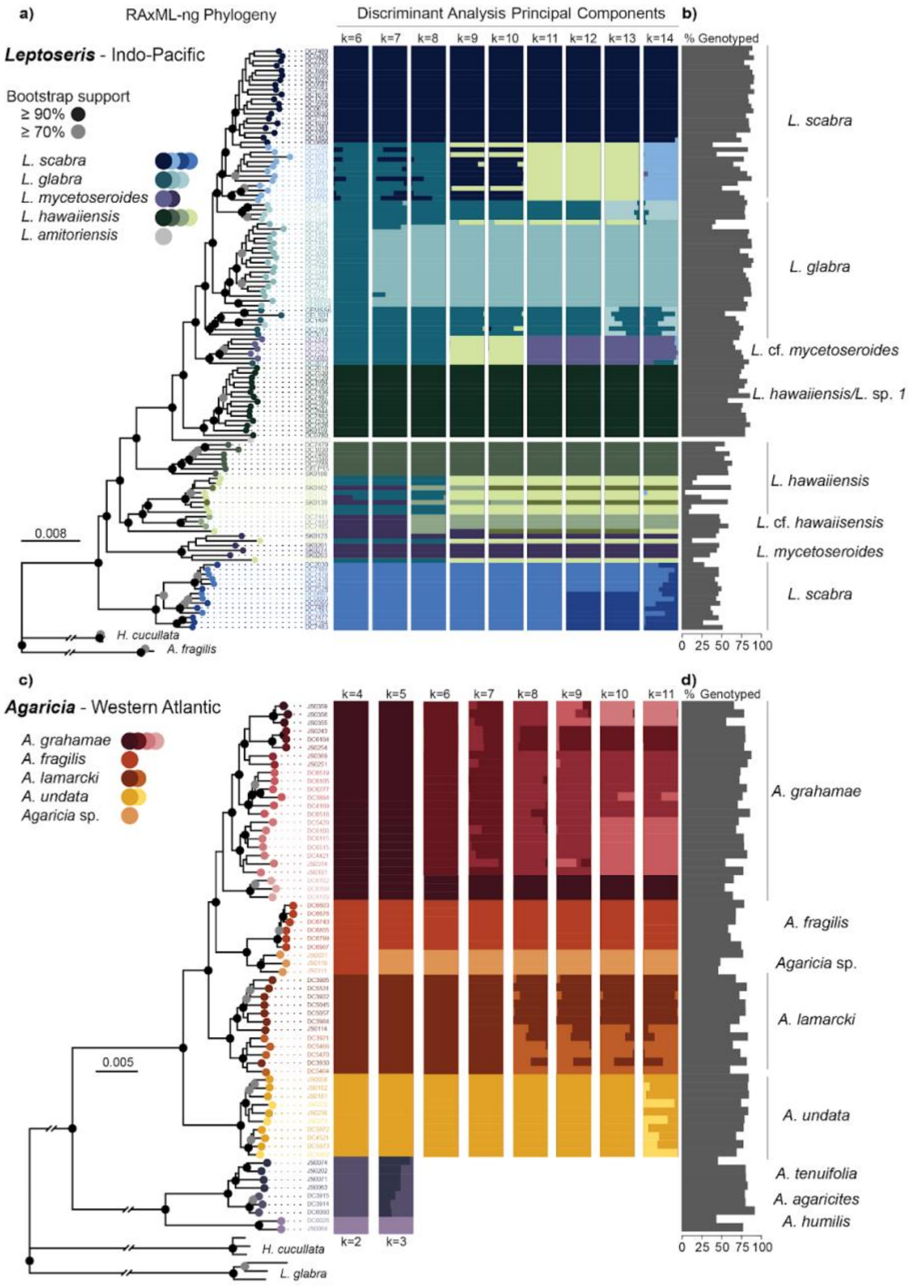


**Supplementary figure S6. *De novo* clustering and ordination methods to assess genetic structure within the genus *Leptoseris* and *Agaricia*.** (a) RAxML-NG tree of the genus *Leptoseris* based on concatenated nextRAD loci, next the posterior membership probabilities of the *de novo* discriminant analysis of principal components (DAPC) based on k=6 to k=14. (b) Barplot representing the percentage of genotyping across *Leptoseris* individuals. (c) RAxML-NG tree of the genus *Agaricia* based on concatenated nextRAD loci, next the posterior membership probabilities of the *de novo* DAPC based on k=6 to k=11, and k=2 to k=3 for the "*Undaria*" group (B) Barplot representing the percentage of genotyping across *Agaricia* individuals.

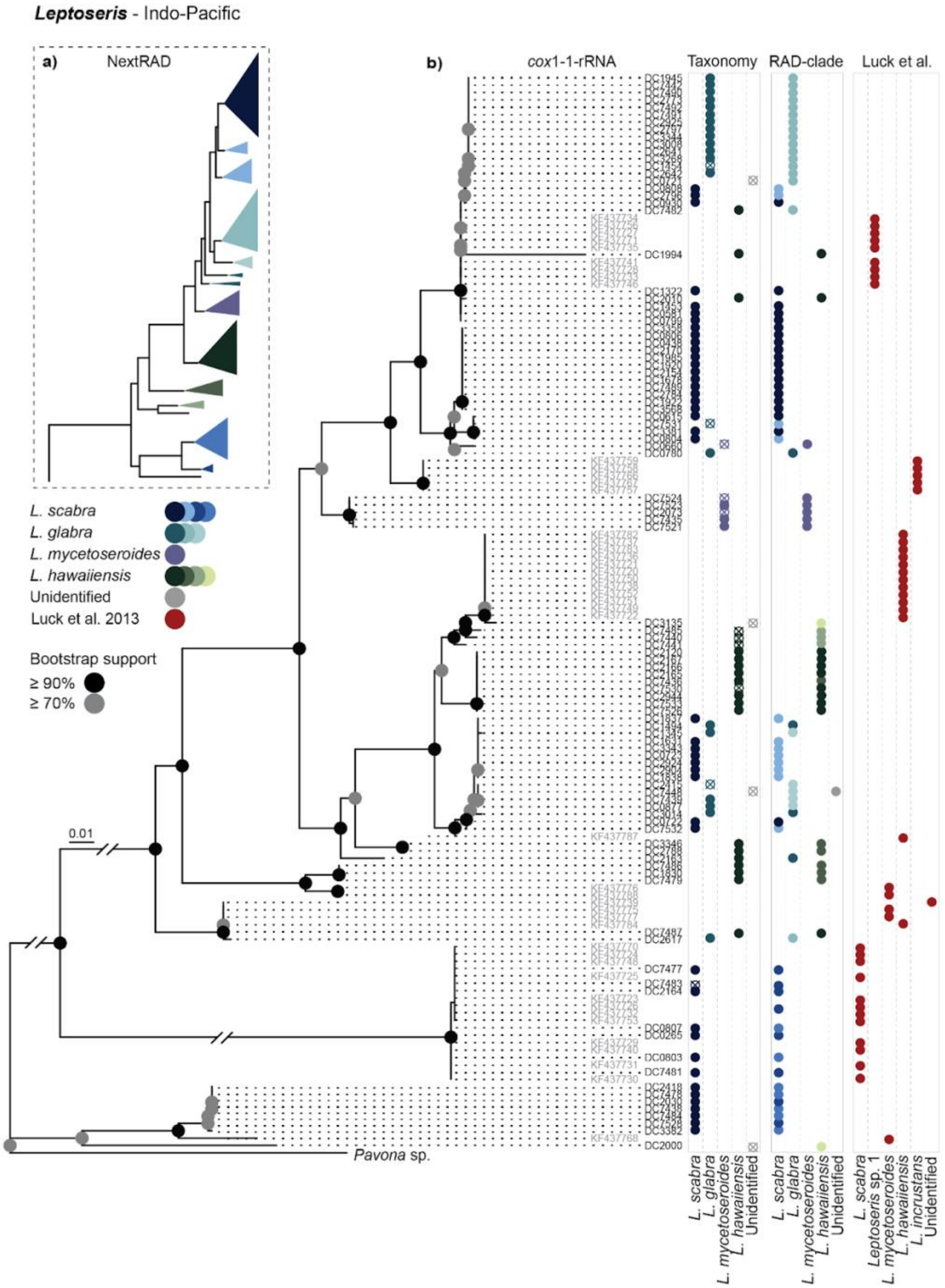


**Supplementary figure S7. Comparison of maximum-likelihood based phylogenies of Australian and Hawaiian *Leptoseris* specimens. (**a) RAxML-NG tree with collapsed nodes of the genus *Leptoseris* based on concatenated nextRAD loci, (b) Mitochondrial *cox*1-1-rRNA marker of a subset (n = 102) of the Australian samples, and Hawaiian samples from Luck *et al.* [17] (grey labels, n = 46), using an individual of *Pavona sp*. as outgroup; next, the taxonomic identification and the nextRAD clade assignment based on the DAPC results.

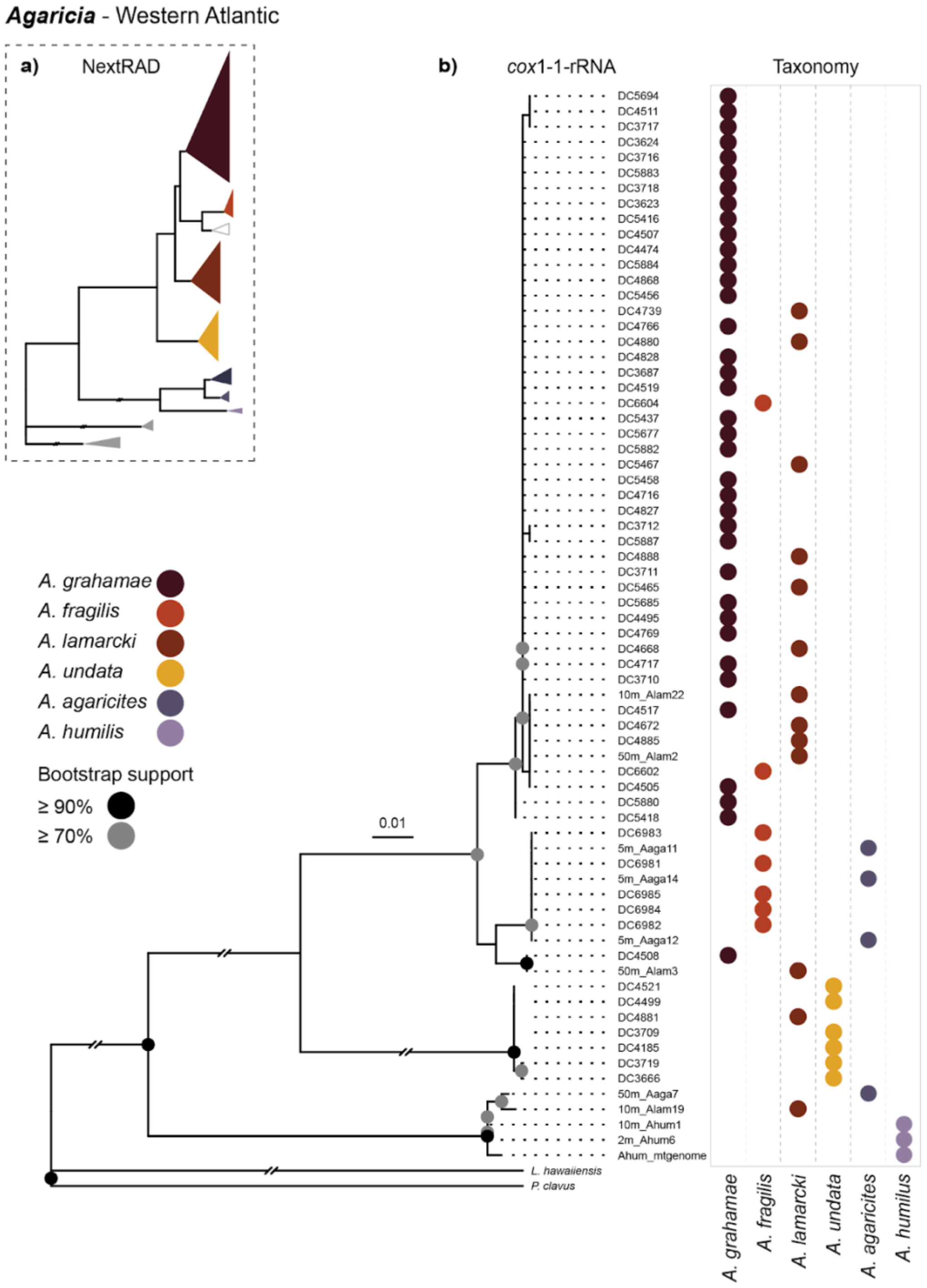


**Supplementary figure S8. Comparison of maximum-likelihood based phylogenies of *Agaricia* specimens.** (a) RAxML-NG tree with collapsed nodes of the genus *Agaricia* based on concatenated nextRAD loci from this study, (b) Mitochondrial *cox*1-1-rRNA marker of *Agaricia* individuals (n = 9) in addition to published *cox*1-1-rRNA sequence data of *Agaricia* individuals (n = 61) from Bongaerts *et al.* (2015) and Medina *et al.* (2006) with *Pavona clavus* (n = 1) and *L. hawaiiensis* (n = 1) as outgroup; next, the taxonomic identification.
